# Age-related differences in encoding-retrieval similarity and their relationship to false memory

**DOI:** 10.1101/2021.07.12.451838

**Authors:** Jordan D. Chamberlain, Caitlin R. Bowman, Nancy A. Dennis

## Abstract

Typical aging is associated with increases in false memory rates among older adults. Such errors are frequently associated with differential neural activity during encoding and retrieval in older compared to younger adults within visual cortices, hippocampus, and front-parietal regions. It remains unknown how pattern similarity reductions relate to false memories in healthy aging. Using encoding-retrieval similarity (ERS) analyses in a sample of younger and older adults, we examined how the similarity of neural patterns between memory phases associated with target and lure objects was impacted by age and contributed to false memory rates. Single-item ERS for targets and lures was reduced by age throughout much of the ventral visual stream and the posterior hippocampus. Furthermore, ERS associated with perceptual lures within the visual stream maintained differential relationships with false memory. Finally, a global ERS metric accounted for age deficits in single-item ERS, but did not contribute to false memory rates. These findings highlight the contribution of age-related reductions in ERS across multiple representational levels to false memories in healthy aging.

## 1. Introduction

Age-related increases in false memories are a significant source of memory impairment in aging (McCabe et al., 2009). Often such errors of commission occur when older adults fail to encode and/or retrieve details from previous experiences, lending to their reliance on gist traces at the time of memory retrieval (Koutstaal, 2003; Koutstaal & Schacter, 1997; Norman & Schacter, 1997; Tun et al., 1998). With respect to perceptual false memories, errors typically occur as a result of increased perceptual similarity between what was viewed during encoding and item lures presented during retrieval (Bowman et al., 2019; Dennis & Turney, 2018; Slotnick & Schacter, 2004). That is, the perceptual similarity between a target and a lure item leads individuals to falsely identify the lure item as ‘old’ during a memory test. Such errors are due in part to increased reliance on perceptual gist signals during retrieval to make the memory decision (i.e., “I have seen a similar object before”), often when more fine-grained perceptual details are unavailable. While the vast majority of false memory research has focused on retrieval processes to explain such memory errors, failures of encoding, as well as the relationship between encoding and retrieval processing, are also key determinants in understanding both memory success and memory errors of commission (Ritchey et al., 2013; Wing et al., 2020).

Within studies of aging, the examination of false memories has largely focused on age differences in blood oxygen level dependent (BOLD) differences between veridical and false memories during memory retrieval. Results of these studies have identified several critical and related findings. Specifically, the studies find that older adults tend to underrecruit occipital regions supporting true memories and overrecruit both prefrontal and lateral temporal regions during the execution of false memories (Dennis, Johnson, et al., 2014; Dennis & Turney, 2018; Paige et al., 2016; Webb & Dennis, 2018). Activity in occipital regions, including early visual cortex and lingual gyrus, are posited to reflect sensory signals that are sensitive to the original presentation of previously viewed items (e.g., targets; Dennis et al., 2012; Slotnick & Schacter, 2004; Wheeler & Buckner, 2004). Greater activity for true compared to false memories in occipital regions has been posited to reflect the fact that such details are available at retrieval for targets, yet not for related lures. Further, past work implicates medial superior frontal gyrus associated with increased monitoring demands in the presence of lure items (Kurkela & Dennis, 2016) and middle and superior temporal gyri associated with retrieval of gist-related memory traces (Gutchess & Schacter, 2012; Webb et al., 2016). This differential recruitment suggests that older adults do not utilize perceptual details from encoding when later faced with the need to make old/new memory decisions during retrieval. Rather, the finding suggest that older adults rely upon the retrieval of gist-based memory traces and top-down monitoring of these gist traces. This conclusion is consistent with behavioral studies that find that, even when older adults encode details that are needed to distinguish between old and new information, they do not take advantage of this information during the retrieval process (Bowman & Dennis, 2015; Bulevich & Thomas, 2012; Cohn et al., 2008; Koutstaal, 2003; Mitchell et al., 2013; Multhaup, 1995; Park et al., 1984; Pezdek, 1987; Rahhal et al., 2002). The foregoing findings further indicate that encoded representations are not be successfully recapitulated during retrieval in the same manner as found in younger adults.

Similar age-deficits in utilizing encoding details to support identification of targets and lure rejection are also observed in research showing that older adults exhibit impairments in the ability to engage in a recall-to-reject retrieval strategy when viewing perceptually similar items, compared with younger adults (Bowman & Dennis, 2015; Castel & Craik, 2003; Gallo, 2006; Jacoby, 1999; Jennings & Jacoby, 1997; Multhaup, 1995). That is, during retrieval, when making memory decisions about targets and perceptually related lures, recall-to-reject theory posits that one must retrieve details regarding past events in order to correctly identify novel information and reject it as new (Gallo, 2006; Jennings & Jacoby, 1997; Multhaup, 1995). Implicit in the recall-to-reject process is that one must have available to them the necessary details from past events in order to carry out the recall-to-reject decision. If item specific details are absent to older adults at the time of retrieval, they rely on the retrieval of gist traces when making memory decisions in the presence of lures (Balota et al., 1999; Kensinger & Schacter, 1999; Koutstaal & Schacter, 1997; Norman & Schacter, 1997; Schacter et al., 1997; Tun et al., 1998) and thus are likely to fail to identify related, yet novel, items as new when the gist traces between the studied item and lure supports the identity of both items.

The notion that neural processes during encoding are recapitulated at retrieval, leading to memory success, has been recently investigated by examining the correlation of neural patterns across memory phases in young adults. That is, using encoding retrieval similarity (ERS) analyses, studies have demonstrated that information shared between two memory phases benefits memory performance (Kuhl et al., 2011; Ritchey et al., 2013; Trelle et al., 2019; Ye et al., 2016). For example, early work by Ritchie and colleagues (2013) showed that the similarity of neural patterns between encoding and retrieval associated with scenes were predictive of successful memory performance in younger adults. Specifically, both single-item and set-level ERS was associated with successful remembering within the ventral visual stream and frontal cortices (Ritchie et al. 2013). This finding is supported a more recent study by Trelle et al. (2020) who also found that ventral temporal cortex ERS was predictive of veridical memory retrieval. Accordingly, research suggests that the similarity in neural patterns within ventral visual cortices across memory phases play a critical role in memory success in younger adults.

In contrast to the positive relationship observed with respect to neural pattern similarity and veridical memories, findings related to neural pattern similarity and false memories are more nuanced. Notably, Chadwick et al. (2016) found that semantic overlap among test items related to increasing pattern similarity in temporal pole, and this semantic-neural overlap contributed to more false memories. Their results suggest neural traces corresponding to semantic relatedness become increasingly similar across trials, resulting in more false memories associated with gist-based processing. Additional work suggests that increased ERS associated with false memories (compared to true memories) is observable in parietal, frontal, and late occipital regions (Lee et al., 2019; Ye et al., 2016). Such findings support computational models of false memory formation such as MINERVA2 that hypothesize that memory traces at retrieval are related to those at encoding (Arndt & Hirshman, 1998). Finally, recent work by Wing et al. (2020) found that the interaction of ERS between visual cortex and hippocampus relates to higher incidents of false memories in younger adults. Taken together, ERS findings highlight how fine-grained information maintained within neural patterns of activity underly erroneous memory performance within younger adults.

Despite these studies, very little work has examined how the recapitulation of brain patterns from encoding to retrieval is affected by age, and whether age-related differences in ERS contribute to age-related increases in false memories. In a recent study, Hill and colleagues (2021) demonstrated that older adults exhibited reduced ERS for veridical scene memory compared to younger adults in select portions of the ventral visual stream, including fusiform cortex. Additionally, the foregoing results suggested that the dedifferentiation of neural patterns during encoding accounted for much of this age reduction in ERS within older adults. Such findings support the idea that older adults do not retain detailed representations of information from encoding to retrieval in the same manner as younger adults, losing the retrieval benefit of perceptual details presented during encoding (Daselaar et al., 2006; Koutstaal, 2003). To this point, reduced distinctiveness of neural representations during both encoding and retrieval has been documented across a handful of studies in the aging literature (Bowman et al., 2019; Koen & Rugg, 2019; Trelle et al., 2019, 2020). For example, Koen & Rugg (2019) showed that neural patterns associated with objects and scenes during encoding were less distinctive (more dedifferentiated) in the parahippocampal place area in older compared to younger adults, and that the distinctiveness of these neural patterns positively predicted subsequent item recognition across the entirety of their sample. While Koen et al. (2019) did not directly assess the relationship between dedifferentiation of neural patterns associated with objects and scenes and false memory performance, the results nonetheless carry important implications for false memories. Most notably, the results suggest that the distinctiveness of neural patterns encoded are related to the ability to recognize items later. Thus, reduced distinctiveness of this information can lead to more liberal memory decisions at retrieval, in particular for information that is similar but not the same as previously studied information. Taken together, this suggests that behavioral and neural components of representational quality decline in older adults and differences in the reactivation of memory representations likely contribute to memory errors in aging.

However, to our knowledge, no study has examined how such encoding-retrieval similarity of neural patterns associated with target and lure items are impacted by age and how age differences in ERS underlie age-related increases in false memories. In order to investigate these points, the current study examined both target-target ERS and well as target-lure ERS throughout cortical regions critical to memory retrieval in both younger and older adults. The study also examined how age differences in ERS correspond to behavioral differences between younger and older adults, with a focus on explaining age-related increases in false memories. We posited that older adults would exhibit age-related reductions in encoding-retrieval similarity, particularly in visual and temporal regions, and that such reductions will be associated with false memory rates in aging. Furthermore, given neural pattern similarity interactions between hippocampus and visual cortex associated with false memories (Wing et al., 2020), we further predicted older adults would demonstrate reduced encoding-retrieval similarity in medial temporal regions, with reductions underlying both veridical and false memories in aging. Finally, given the differential role of parietal and frontal regions during encoding and retrieval, we did not expect age-related reductions in those regions, but expected them to relate to false memory performance.

## 2. Materials and Methods

### 2.1 Participants

27 young adults and 27 older adults were recruited from The Pennsylvania State University and State College community, respectively. All participants were right-handed, native English speakers who were screened for contraindications for MRI and neurological and psychiatric disorders. All received monetary compensation for their participation. Two young adults were excluded from analyses due to failure to complete the task (1 subject) and movement in excess of 2.5 mm within a run (1 subject), leaving data from 25 young adults included in all analyses (M = 22.32 years old, SD = 3.275 years, range = 18-31; 17 females). Four older adults were excluded from analyses due to technical issues during data collection (2 subjects) and movement in excess of 2.5 mm within a run (2 subjects), leaving data from 23 older adults included in all analyses (M = 71.04 years old, SD = 8.04 years old, range = 60-84; 12 females). Other univariate analyses from the young adult group (Bowman et al., 2017; Bowman & Dennis, 2016) and multi-voxel pattern analyses from retrieval (Bowman et al., 2019) have been reported previously. All participants completed written informed consent and all experimental procedures were approved by Penn State’s Office of Research Protections. Additionally, all older adults showed no indicators of cognitive impairment, assessed by the Mini Mental State Exam (Folstein et al., 1983) scores > 28 (mean = 29.65 (0.65) or depression, assessed by the Geriatric Depression Scale ((Sheikh & Yesavage, 1986) (mean = 0.88 (1.08)).

### 2.2 Stimuli

The stimuli description and task procedures were detailed in our previously published papers (Bowman et al., 2019; Bowman & Dennis, 2016). They are reprinted here for completeness. Stimuli consisted of 316 images of common objects collected from the Band of Standardized Stimuli database (Brodeur et al., 2010) as well as an Internet image search. Images were cropped and resized to approximately 400 × 400 pixels and equated for resolution. Images were displayed at a screen resolution of 1024 (H) × 768 (V) at 75 Hz. At the viewing distance of 143 cm, the display area was 20° (H) ° 16° (V) with experimental stimuli subtending 5° (H) × 4° (V).

Encoding items included 96 images, each an exemplar from a distinct category or conceptual theme. At retrieval, four types of items were presented: targets, item lures, thematic lures, and novel lures (see Figure 1 for examples). Item lures were an alternative exemplar of the same item presented at encoding. Thematic lures were a new item within the same general category or theme of an item presented at encoding. Novel lures were items drawn from categories not presented during encoding. (See Bowman et al., 2019 for a full description of stimuli norming).

**Figure 1.**
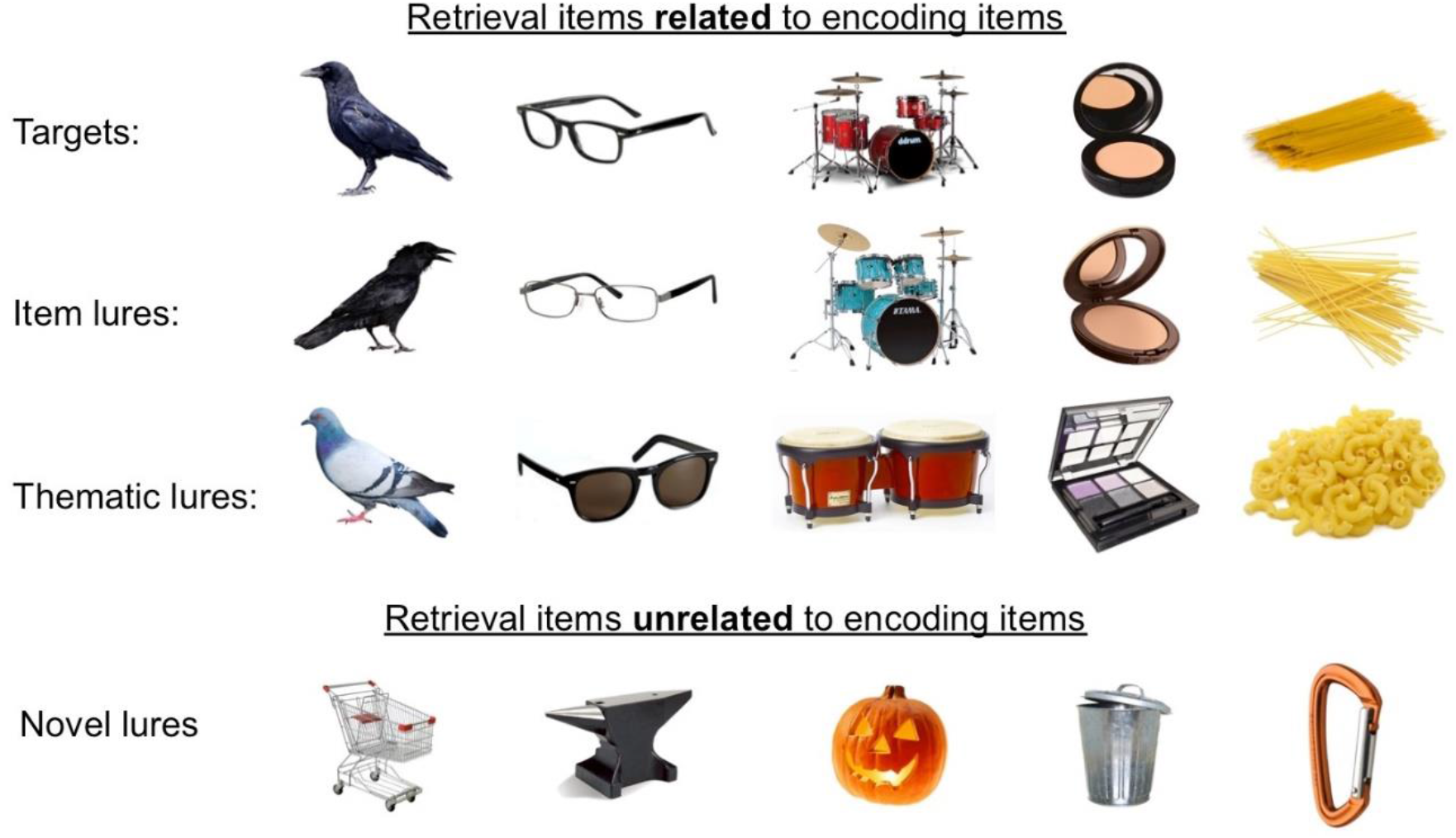
Example Stimuli. Participants viewed objects during encoding while making size judgements. During retrieval, participants viewed objects that were targets (the same object viewed during encoding), item lures (perceptually related objects), thematic lures (object from the same category), and novel lures (completely unrelated objects). Figure reproduced from Bowman et al., 2019.

### 2.3 Experimental Design

Participants completed the encoding and retrieval phases in a single scanning session. Stimuli were displayed using COGENT in MATLAB (MathWorks, Natick, MA) and were back-projected onto a screen that participants viewed through a mirror attached to the head coil. Behavioral responses were recorded using a four-button response box. Scanner noise was reduced with headphones, and cushioning was used to reduce head motion.

Encoding was incidental, and participants were instructed to make a size judgment for each of 96 items (i.e., “Is this item bigger or smaller than a shoebox?”). Each image was presented for 1500 msec followed by 500-msec additional responding time before automatically advancing to the next item. Each item was followed by a variable intertrial fixation (*M* = 2470 msec, *SD* = 1760 msec) to aid in deconvolving the hemodynamic response (Dale, 1999). After the encoding phase, participants underwent structural scans and were given instructions for retrieval. During retrieval, participants were presented with all 96 studied items (targets), both the item and thematic lures from each of the encoding categories (96 of each), and 28 novel lures. All stimulus categories were presented in an intermixed fashion during each run, pseudorandomly ordered to ensure that no more than three stimuli of the same type were presented sequentially. Each stimulus was displayed for 3 sec while participants made their memory responses. Each trial was followed by a variable intertrial interval (M = 2340 msec, SD = 1440 msec). In accord with typical instructions for the remember–know–new paradigm, participants were instructed to respond “remember” if they believed that the item was presented during the first phase of the experiment and they remembered specific, vivid details of its prior presentation. Participants were instructed to respond “familiar” if they believed the item was presented during the first phase but they could not recall specific details about its prior presentation. Instead of the typical “new” response, participants were asked to respond with two distinct “new” options— “unfamiliar” or “different”(Matzen et al., 2011). Participants were instructed to respond “different” if they believed the item was new and they could recall aspects of truly presented item(s) that indicated the item was not presented during the study phase (i.e., recollection rejection). Participants were instructed to respond “unfamiliar” if they believed an item was new because it did not resemble or bring to mind anything from the previous encoding phase (i.e., unfamiliar item). A practice session before the retrieval scans ensured that participants understood the retrieval instructions and the distinction between each response option before beginning the retrieval scans.

### 2.4 fMRI data acquisition

Imaging data was acquired using a Siemens (Berlin, Germany) 3-T scanner equipped with a 12-channel head coil. A T1-weighted sagittal localizer was acquired to locate the AC–PC. Images were then prescribed at an oblique angle that prevented data collection in the area of the orbits. A high-resolution anatomical image was acquired in the sagittal plane using a magnetization prepared rapid gradient echo sequence with a 1650-msec repetition time, 2.03-msec echo time, 256-mm field of view, 256-mm2 matrix, 160 sagittal slices, and 1-mm slice thickness for each participant. Echoplanar functional images were acquired using a descending acquisition, 2500-msec repetition time, 25-msec echo time, 240-mm field of view, an 80-mm2 matrix, and 42 axial slices with 3-mm slice thickness, resulting in 3-mm isotropic voxels. Ninety-one volumes were collected in each of two functional runs of the encoding task. One hundred seventy-five volumes were collected in each of the four functional runs of the retrieval task.

### 2.5 Statistical analysis

#### Behavioral data

We first examined overall memory performance (d’) by collapsing scores across lure types and old (‘remember’, ‘familiar’) and new (‘different’, ‘unfamiliar’) responses. We used planned *t* tests to examine age differences in d’. We were also interested in whether age differences were driven by hits or false alarms. To do so, we conducted a mixed-factor ANOVA with age and trial type as factors and proportion of items labeled ‘old’ as the dependent variable.

#### 2.5.1 fMRI pre-processing

Anatomical and functional images were skull stripped using the Brain Extraction Tool (BET) in FSL version 5.0.9 (www.fmrib.ox.ac.uk/fsl). FSL’s MCFLIRT function was then used for within-run realignment and motion correction in each functional run, aligning all volumes to the middle volume. Between-run realignment was performed using Advanced Normalization Tools (ANTs: http://stnava.github.io/ANTs/): the first volume of each run was aligned to the first volume of the first run (i.e., the first volume from the first encoding run), and that transformation was then applied to all volumes in the corresponding run. These realigned functional images were then submitted to FSL’s fMRI Expert Analysis Tool (FEAT) where they were high-pass filtered at 100 seconds and underwent minimal spatial smoothing at 2mm FWHM.

#### 2.5.2 Regions of Interest

To define subject-specific anatomical regions of interest (ROIs) in native space, we submitted the high-resolution anatomical image from each subject to Freesurfer version 6 (https://surfer.nmr.mgh.harvard.edu/). Based on our interest in the correspondence of neural patterns between encoding and retrieval related to visual memories, we selected the following anatomical ROIs throughout the ventral visual stream: early occipital cortex (combination of lingual gyrus and cuneus labels in Freesurfer), lateral occipital cortex, posterior fusiform cortex (defined as the portion of fusiform cortex posterior to the most anterior slice of parahippocampal cortex), and inferotemporal cortex. Regions within the fronto-parietal retrieval network and MTL were also selected based on their involvement in veridical and false memory processing (Devitt & Schacter, 2016; Kurkela & Dennis, 2016): superior frontal gyrus, superior parietal cortex, precuneus, and angular gyrus. Among MTL regions we selected entorhinal cortex, anterior and posterior hippocampus. To define the anterior and posterior hippocampal regions we split each participants hippocampus at the middle slice. If a participant had an odd number of hippocampal slices, the middle slice was assigned to the posterior hippocampus (Bowman & Zeithamova, 2018). Finally, based on our prior work in the area of false memories and gist processing (Webb et al., 2016), we also included the temporal pole. Each ROI was broken down bilaterally based on recent work suggesting that laterality is critical to assessing the relevance of neural patterns associated with false memories (Dennis et al., 2012; Garoff-Eaton et al., 2006; Koutstaal et al., 2001; Slotnick & Schacter, 2006; Wing et al., 2020). We collapsed across hemispheres for regions that failed to exhibit a significant age X trial type X hemisphere interaction for subsequent individual differences analyses (see Results-*Individual Differences between Trial Type ERS and False Memories)*.

#### 2.5.3 Multivariate pattern analyses

To estimate neural patterns associated with individual trials, a GLM was generated using FSL’s FEAT with one regressor for each trial at both encoding and retrieval (48 trial regressors for encoding runs, 79 for each retrieval run). An additional six nuisance regressors were included in each run corresponding to translational and rotational movement. From these models, individual whole-brain beta parameter maps were generated for each trial at encoding and retrieval. In a given parameter map, the value in each voxel represented the regression coefficient for that trial’s regressor in a multiple regression containing all other trials in the run and the motion parameters. These beta parameter maps were then concatenated across runs and submitted to the CoSMoMVPA toolbox (https://www.cosmomvpa.org/) in Matlab version 2017b (https://www.mathworks.com/) for ERS analyses.

#### 2.5.4 Encoding-retrieval pattern similarity (ERS)

We performed an ERS analysis based on retrieval trial type (target; item lure; theme lure) to understand how aging affects the recapitulation of neural patterns between encoding and retrieval related to the viewing of targets and related lures. By assessing ERS values based on the retrieval trial type, we can assess how the correspondence of information between encoding and retrieval underlies false memory performance, and how this correspondence is impacted by processes of healthy aging. To characterize how the correspondence of neural patterns from encoding to retrieval is related to different trial types and how that differs with age, we used a representational similarity analysis (Kriegeskorte et al., 2008) in which we calculated the similarity score (Pearson’s r; Ritchey et al., 2013) between the beta parameter pattern of a given item during encoding and that of 1) its re-representation as a target during retrieval, 2) the presentation of its corresponding item lure during retrieval, and 3) the presentation of its corresponding thematic lure during retrieval. After all of these r-values were computed for each of 96 studied items, a Fisher z-transformation was performed and values within a condition (i.e., target-target, target-item lure, target-theme lure; Fig 2A) were averaged within each subject and divided by their standard deviation, generating an ERS effect size for each condition (Bowman & Zeithamova, 2018). This process resulted in three ERS values per subject, per ROI, corresponding to our three conditions of interest (target-target ERS; target-item lure ERS; and target-thematic lure ERS). Trials corresponding to novel lures were estimated in the single trial model but were not included in initial ERS analyses as novel lures at retrieval did not have corresponding items during encoding. Novel lures were later used in the calculation of global ERS. To test for differences across retrieval trial types in the correspondence of patterns between encoding and retrieval, we computed a 2 (hemisphere: left, right) × 2 (age group: young, older) × 3 (ERS comparison: target-target, target-item lure, target-thematic lure) mixed-factors ANOVA per ROI.

**Figure 2.**
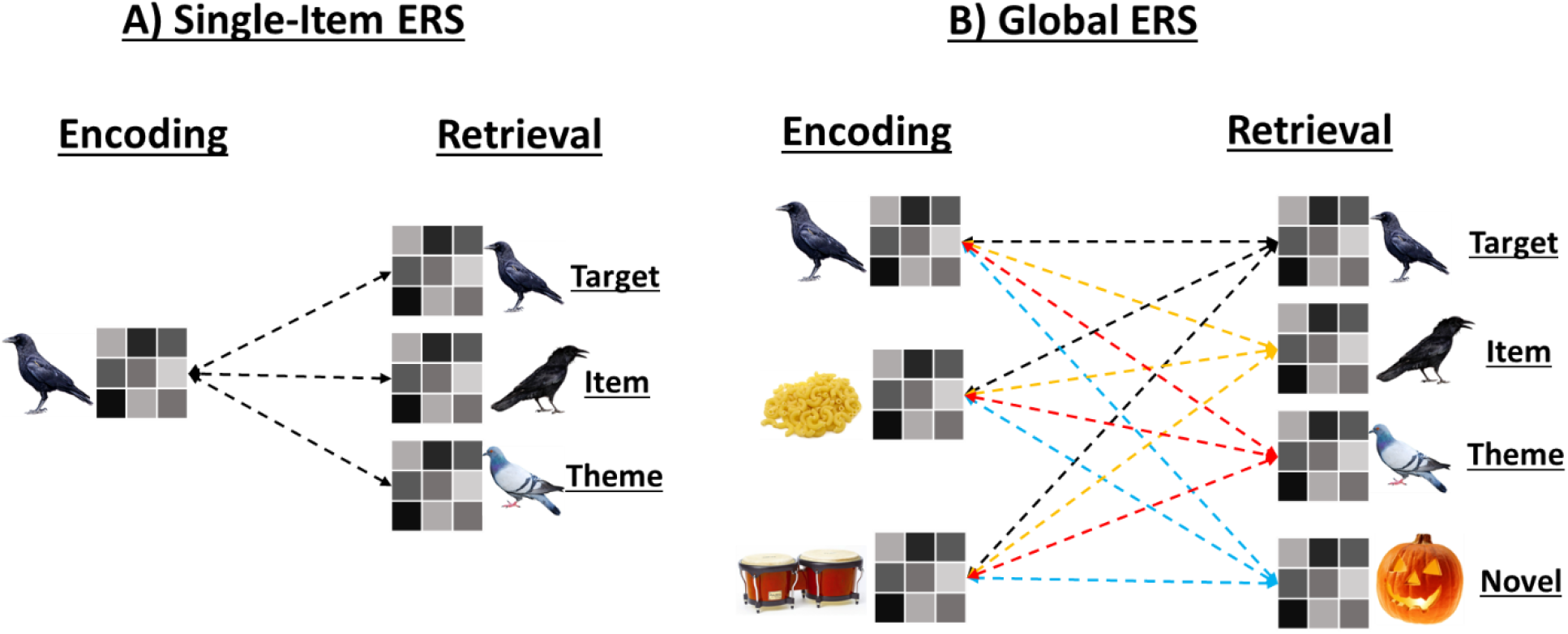
Schematic of ERS metrics. 2A) We calculated ERS between single items at encoding and their corresponding target, item lure, and thematic lure at retrieval. 2B) We created a global ERS metric per ROI in which we calculated the similarity of each trial at encoding with each trial at retrieval to account for baseline group differences in neural similarity.

**Figure 3.**
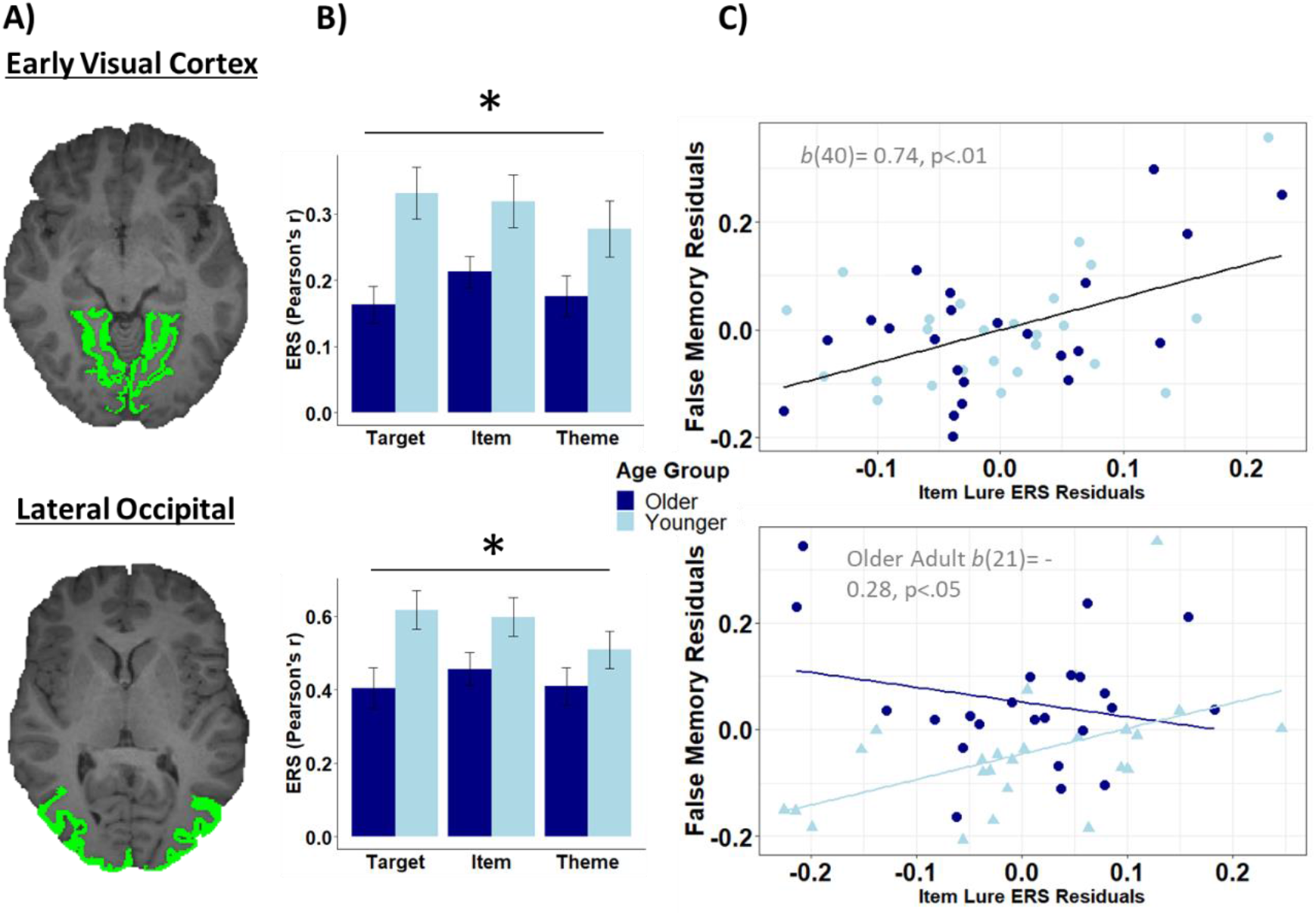
Visual cortex ERS and relationships with behavior. 3A) Early visual and lateral occipital cortex ROIs depicted in a representative participant. 3B) We observed age deficits in ERS throughout portions the ventral visual stream. * indicates age effect *p* < .05. 3C) Item Lure ERS positively predicted false memory rates regardless of age in early visual cortex, while the relationship between item lure ERS and false memory rates was moderated by age in lateral occipital cortex.

To assess how the recapitulation of brain patterns relates to behavioral performance, we computed separate multiple regression models for each ROI with overall behavioral false alarm rates as the dependent measure. We computed the foregoing multiple regression models using the following predictors simultaneously: 1) target-target, target-item lure, and target-thematic lure ERS values 2) age group (young, older) as a categorical predictor, and 3) age × ERS value interaction effects for each of the ERS comparisons.

Recent research suggests that age differences in neural recapitulation can be accounted for by baseline differences in neural similarity between younger and older adults, as older adults have reduced pattern similarity compared to younger adults (Hill et al., 2021). To account for the possibility of such baseline age differences in the current study, we calculated a global ERS value within each region of interest to estimate the general degree of recapitulation between items viewed during encoding and retrieval (i.e., the similarity of all items at retrieval and all items at encoding). Recent work by Ye et al. (2016) and Wing et al. (2020) have found global ERS to be associated with false memory rates in younger adults. Furthermore, such a global ERS metric is posited to reflect the general strength of recapitulation according to global matching models of episodic memory (Arndt & Hirshman, 1998). It is therefore possible that older adults have a reduced baseline ERS as compared to younger adults, and this global ERS explains variance above and beyond single-item ERS when predicting false memory rates. To account for this possibility, we calculated the similarity score between a beta parameter of a given trial at retrieval and that of every item during encoding within each region of interest (Fig 2B). We replicated the above analyses for regions that exhibited a significant age effect while including this global ERS metric as a covariate.

Finally, to correct for multiple comparisons and reduce the effect of false positives, we conducted permutation testing for each significant finding. Specifically, we shuffled the data for every participant and reconducted the ANOVA or multiple linear regression model (Kherad-Pajouh & Renaud, 2015). We repeated this procedure 10,000 times per statistical test to build a null distribution to simulate the potential significance that could be obtained if our effects did not exist.

## 3. Results

### 3.1 Behavioral Results

Behavioral results from this dataset are reported in Bowman et al. (2019), we report the major findings here for completeness. Overall memory performance was indexed by d’ collapsed across lure types and across multiple old (‘remember’, ‘familiar’) and new (‘different’, ‘unfamiliar’) responses. As expected, young adults showed significantly higher d’ than older adults [t(46) = 3.10, p = 0.003], indicating that older adults did not distinguish between old and new items during retrieval as well as young adults.

We were also interested in whether this overall age deficit was driven by an age-related increase in false alarms. We therefore computed a mixed-factor ANOVA with age and trial type as factors and proportion of items labeled ‘old’ as the dependent variable. Results revealed a significant main effect of trial type [*F*(1.83,84.15) = 708.94, *p* < 0.001] with the highest proportion of endorsement for targets (*M* = 0.81, *SD* = 0.10), followed by item lures (*M* = 0.41, *SD* = 0.19), then thematic lures (*M* = 0.10, SD = 0.10), and the lowest endorsement of novel lures (*M* = 0.02, *SD* = 0.03). There was also a significant main effect of age [*F*(1,46) = 12.66, *p* = 0.001], with older adults endorsing items as old at higher rates (*M* = 0.38, *SD* = 0.09) than young adults (*M* = 0.29, *SD* = 0.09). Finally, there was a significant age × trial type interaction [*F*(1.83,84.15) = 17.80, *p* < 0.001]. Post-hoc independent samples *t*-tests showed that older adults had more false recognitions than young adults for item lures [*t*(46) = 4.72, *p* < 0.001] and thematic lures [*t*(46) = 2.32, *p* = 0.03] but that reliable age differences in ‘old’ responses did not emerge for targets [*t*(46) = 1.01, *p* > 0.05] or novel lures [*t*(46) = 0.63, *p* > 0.05].

### 3.2 Encoding-Retrieval Similarity (ERS) Trial Type Differences

We next examined age and trial differences among ERS values in each region of interest. To do so we conducted a 2(hemisphere: left, right) × 3(trial type: target, item lure, theme lure) × 2(age group: young, old) within-between repeated measures ANOVA within each region of interest. See Table 1 for complete results.

**Table 1.**
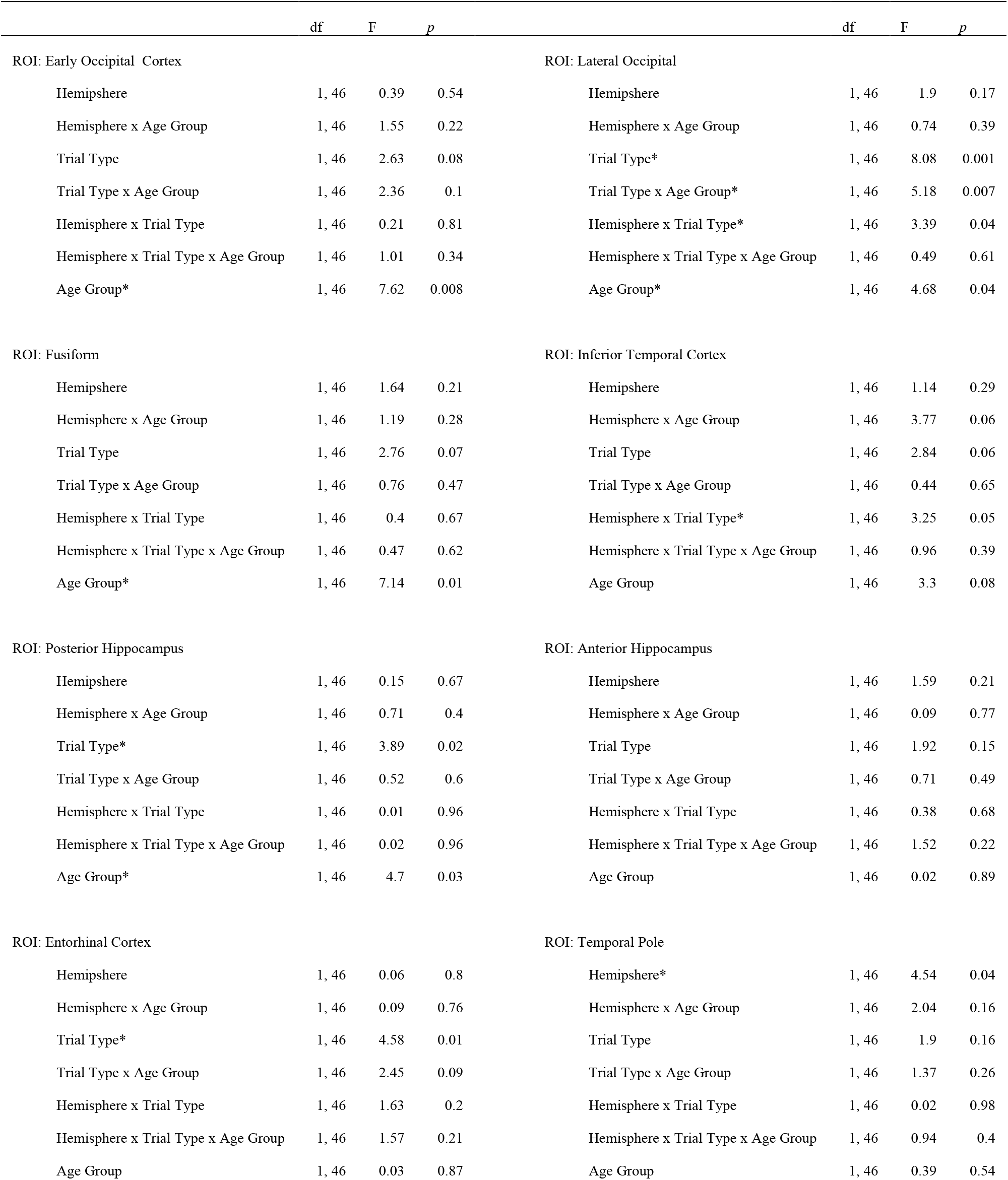
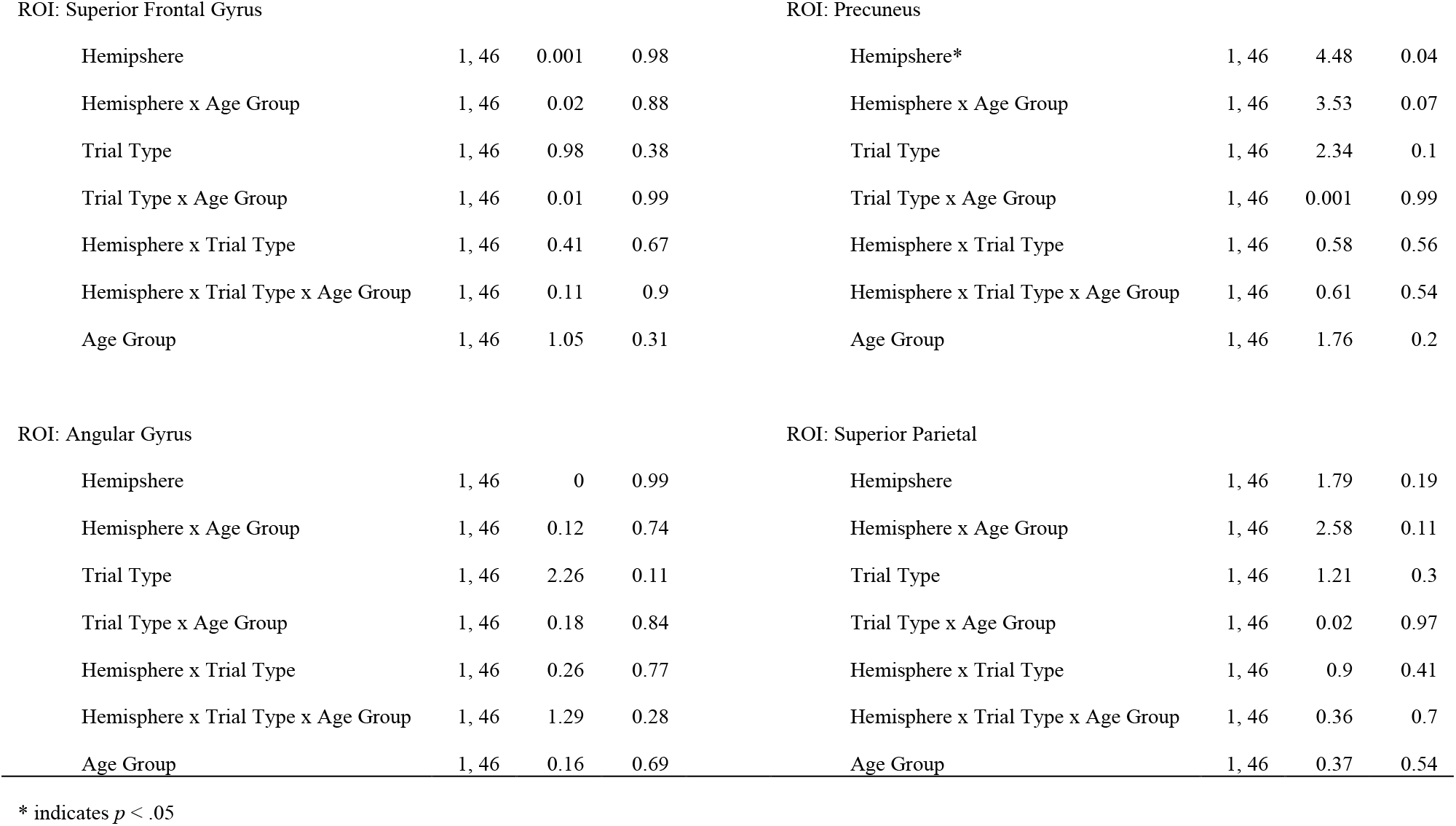
Results for the 2 (hemisphere: left, right) × 2 (age group: young, older) × 3 (ERS comparison: target-target, target-item lure, target-thematic lure) mixed-factors ANOVA per ROI.

|  | df | F | p |  | df | F | p |
| --- | --- | --- | --- | --- | --- | --- | --- |
| ROI: Early Occipital Cortex |  |  |  | ROI: Lateral Occipital |  |  |  |
| Hemisphere | 1, 46 | 0.39 | 0.54 | Hemisphere | 1, 46 | 1.9 | 0.17 |
| Hemisphere x Age Group | 1, 46 | 1.55 | 0.22 | Hemisphere x Age Group | 1, 46 | 0.74 | 0.39 |
| Trial Type | 1, 46 | 2.63 | 0.08 | Trial Type* | 1, 46 | 8.08 | 0.001 |
| Trial Type x Age Group | 1, 46 | 2.36 | 0.1 | Trial Type x Age Group* | 1, 46 | 5.18 | 0.007 |
| Hemisphere x Trial Type | 1, 46 | 0.21 | 0.81 | Hemisphere x Trial Type* | 1, 46 | 3.39 | 0.04 |
| Hemisphere x Trial Type x Age Group | 1, 46 | 1.01 | 0.34 | Hemisphere x Trial Type x Age Group | 1, 46 | 0.49 | 0.61 |
| Age Group* | 1, 46 | 7.62 | 0.008 | Age Group* | 1, 46 | 4.68 | 0.04 |
| ROI: Fusiform |  |  |  | ROI: Inferior Temporal Cortex |  |  |  |
| Hemisphere | 1, 46 | 1.64 | 0.21 | Hemisphere | 1, 46 | 1.14 | 0.29 |
| Hemisphere x Age Group | 1, 46 | 1.19 | 0.28 | Hemisphere x Age Group | 1, 46 | 3.77 | 0.06 |
| Trial Type | 1, 46 | 2.76 | 0.07 | Trial Type | 1, 46 | 2.84 | 0.06 |
| Trial Type x Age Group | 1, 46 | 0.76 | 0.47 | Trial Type x Age Group | 1, 46 | 0.44 | 0.65 |
| Hemisphere x Trial Type | 1, 46 | 0.4 | 0.67 | Hemisphere x Trial Type* | 1, 46 | 3.25 | 0.05 |
| Hemisphere x Trial Type x Age Group | 1, 46 | 0.47 | 0.62 | Hemisphere x Trial Type x Age Group | 1, 46 | 0.96 | 0.39 |
| Age Group* | 1, 46 | 7.14 | 0.01 | Age Group | 1, 46 | 3.3 | 0.08 |
| ROI: Posterior Hippocampus |  |  |  | ROI: Anterior Hippocampus |  |  |  |
| Hemisphere | 1, 46 | 0.15 | 0.67 | Hemisphere | 1, 46 | 1.59 | 0.21 |
| Hemisphere x Age Group | 1, 46 | 0.71 | 0.4 | Hemisphere x Age Group | 1, 46 | 0.09 | 0.77 |
| Trial Type* | 1, 46 | 3.89 | 0.02 | Trial Type | 1, 46 | 1.92 | 0.15 |
| Trial Type x Age Group | 1, 46 | 0.52 | 0.6 | Trial Type x Age Group | 1, 46 | 0.71 | 0.49 |
| Hemisphere x Trial Type | 1, 46 | 0.01 | 0.96 | Hemisphere x Trial Type | 1, 46 | 0.38 | 0.68 |
| Hemisphere x Trial Type x Age Group | 1, 46 | 0.02 | 0.96 | Hemisphere x Trial Type x Age Group | 1, 46 | 1.52 | 0.22 |
| Age Group* | 1, 46 | 4.7 | 0.03 | Age Group | 1, 46 | 0.02 | 0.89 |
| ROI: Entorhinal Cortex |  |  |  | ROI: Temporal Pole |  |  |  |
| Hemisphere | 1, 46 | 0.06 | 0.8 | Hemisphere* | 1, 46 | 4.54 | 0.04 |
| Hemisphere x Age Group | 1, 46 | 0.09 | 0.76 | Hemisphere x Age Group | 1, 46 | 2.04 | 0.16 |
| Trial Type* | 1, 46 | 4.58 | 0.01 | Trial Type | 1, 46 | 1.9 | 0.16 |
| Trial Type x Age Group | 1, 46 | 2.45 | 0.09 | Trial Type x Age Group | 1, 46 | 1.37 | 0.26 |
| Hemisphere x Trial Type | 1, 46 | 1.63 | 0.2 | Hemisphere x Trial Type | 1, 46 | 0.02 | 0.98 |
| Hemisphere x Trial Type x Age Group | 1, 46 | 1.57 | 0.21 | Hemisphere x Trial Type x Age Group | 1, 46 | 0.94 | 0.4 |
| Age Group | 1, 46 | 0.03 | 0.87 | Age Group | 1, 46 | 0.39 | 0.54 |

ROI: Superior Frontal Gyrus
|  |  |  |  |
| --- | --- | --- | --- |
| Hemisphere | 1, 46 | 0.001 | 0.98 |
| Hemisphere x Age Group | 1, 46 | 0.02 | 0.88 |
| Trial Type | 1, 46 | 0.98 | 0.38 |
| Trial Type x Age Group | 1, 46 | 0.01 | 0.99 |
| Hemisphere x Trial Type | 1, 46 | 0.41 | 0.67 |
| Hemisphere x Trial Type x Age Group | 1, 46 | 0.11 | 0.9 |
| Age Group | 1, 46 | 1.05 | 0.31 |

ROI: Precuneus
|  |  |  |  |
| --- | --- | --- | --- |
| Hemisphere* | 1, 46 | 4.48 | 0.04 |
| Hemisphere x Age Group | 1, 46 | 3.53 | 0.07 |
| Trial Type | 1, 46 | 2.34 | 0.1 |
| Trial Type x Age Group | 1, 46 | 0.001 | 0.99 |
| Hemisphere x Trial Type | 1, 46 | 0.58 | 0.56 |
| Hemisphere x Trial Type x Age Group | 1, 46 | 0.61 | 0.54 |
| Age Group | 1, 46 | 1.76 | 0.2 |

ROI: Angular Gyrus
|  |  |  |  |
| --- | --- | --- | --- |
| Hemisphere | 1, 46 | 0 | 0.99 |
| Hemisphere x Age Group | 1, 46 | 0.12 | 0.74 |
| Trial Type | 1, 46 | 2.26 | 0.11 |
| Trial Type x Age Group | 1, 46 | 0.18 | 0.84 |
| Hemisphere x Trial Type | 1, 46 | 0.26 | 0.77 |
| Hemisphere x Trial Type x Age Group | 1, 46 | 1.29 | 0.28 |
| Age Group | 1, 46 | 0.16 | 0.69 |

ROI: Superior Parietal
|  |  |  |  |
| --- | --- | --- | --- |
| Hemisphere | 1, 46 | 1.79 | 0.19 |
| Hemisphere x Age Group | 1, 46 | 2.58 | 0.11 |
| Trial Type | 1, 46 | 1.21 | 0.3 |
| Trial Type x Age Group | 1, 46 | 0.02 | 0.97 |
| Hemisphere x Trial Type | 1, 46 | 0.9 | 0.41 |
| Hemisphere x Trial Type x Age Group | 1, 46 | 0.36 | 0.7 |
| Age Group | 1, 46 | 0.37 | 0.54 |

\* indicates $p < .05$

Largely in line with our predictions, we observed a main effect of age within the ventral visual cortex, including early visual cortex [*F*(1,46) = 7.62 (*p*=.008)], lateral occipital cortex [*F*(1,46) = 4.67 (*p*=.04)], fusiform gyrus [*F*(1,46) = 7.14 (*p*=.01)], and posterior hippocampus [*F*(1,46) = 4.70 (*p* =.03)], with all regions exhibiting lower overall ERS in older compared to younger adults. However, unexpectedly, a main effect of hemisphere was found in temporal pole [*F*(1,1) = 4.54 (*p* =.04)] and precuneus [*F*(1,1) = 4.47 (*p* =.04)], driven by greater ERS in the left than right hemisphere of the temporal pole (t(47) = 2.17, *p* =.03) and greater ERS in the right than left hemisphere of the precuneus (*t*(47) = −2.14, *p* =.04). Also unexpectedly, main effects of trial type also emerged, with three different patterns across regions. First, ERS was higher for targets and items lures compared to thematic lures in lateral occipital cortex. Second, ERS was higher for item lures compared to thematic lures in the posterior hippocampus while there no differences between targets and lures of either type. Third, ERS was higher for item lures than targets and thematic lures in entorhinal cortex.

In line with our overall predictions, there was a significant hemisphere × trial type interaction effect in lateral occipital cortex [*F*(1,46) = 3.39 (*p* =.04)] and inferior temporal cortex [*F*(1,46) = 3.25 (*p* =.05)]. In lateral occipital cortex, the interaction was driven by greater ERS for targets than thematic lures in the right (*t*(47)= 3.48, *p* =.001), but not left (*t*(47)= 1.68, *p* =.10), hemisphere. In the inferior temporal cortex, the interaction was driven by greater ERS for item lures and targets compared to thematic lures in the right hemisphere (*p*’s <.05), but not in left hemisphere (*p*’s >.05). Finally, lateral occipital cortex also exhibited a significant trial type X age group interaction *F*(1,2) = 5.18 (*p* =.007), driven by greater ERS for targets, and item lures, in younger compared to older adults (*p*’s<.05).

Since it is possible that age effects in ERS can be accounted for by a general reduction in neural similarity in older compared to younger adults (Hill et al., 2021), we investigated the effect of global ERS on age differences in single-item level ERS. To do so we repeated the above 2(hemisphere: left, right) × 3(trial type: target, item lure, theme lure) × 2(age group: young, old) within-between repeated measures ANOVA while including global ERS as a covariate. When including the global ERS as a covariate in the analysis, the main effect of age in early visual cortex, lateral occipital cortex, fusiform gyrus, and posterior hippocampus became nonsignificant (all *p’s* > .05). The only significant age interaction to remain was the trial type X age group interaction in lateral occipital cortex *F*(1,2) = 5.18 (*p* =.007) reflecting greater ERS for targets and item lures in younger compared to younger adults.

To confirm significant findings and correct for the effect of false positives, we reconducted the above analyses while conducting permutation testing. Specifically, we repeatedly shuffled the data randomly and reconducted our ANOVAs (Kherad-Pajouh & Renaud, 2015). We repeated this procedure 10,000 times for each ANOVA to build a null distribution to simulate the potential difference in means that could be obtained if our effects did not exist. We found no differences in significance after conducting permutation testing.

### 3.3 Individual Differences between Trial Type ERS and False Memories

To examine how ERS of targets and lures relates to false memories, for each region of interest we conducted a multiple linear regression in which the outcome was false memory rate, and predictors included target ERS, item lure ERS, thematic lure ERS, and their interactions with age group (see Table 2). As we did not observe any significant age X trial type X hemisphere interactions in the previous analysis, we collapsed ERS values across hemispheres. In early visual cortex, item lure ERS positively predicted false memory rates (b(40)= 0.74, *p*=.005), while thematic lure ERS negatively predicted false memory rate (b(40)= −0.50, *p*=.04). In lateral occipital cortex, we observed a significant age group X item lure ERS interaction (b(40) = 0.71, *p*=.03), such that item lure ERS negatively predicted false memory rates in older adults (b(21) = −0.29, *p*<.05) but not younger adults (b(23) = 0.04, *p* >.05). Finally, in posterior hippocampus, thematic lure ERS negatively predicted false memory rates (b(40) = −0.70, *p*=.04).

**Table 2.** Relationship between ERS and false memory rates with and without controlling for global ERS.

| Predictor | Beta | t | p | Predictor | Beta | t | p |
| --- | --- | --- | --- | --- | --- | --- | --- |
| ROI: Early Occipital Cortex |  |  |  | ROI: Early Occipital Cortex |  |  |  |
| Intercept* | 0.25 | 5.17 | <.001 | Intercept* | 0.25 | 5.12 | <.001 |
| Age | -0.1 | -1.54 | 0.13 | Age | -0.01 | -1.47 | 0.15 |
| Target ERS | -0.18 | -0.68 | 0.5 | Target ERS | -0.17 | -0.62 | 0.54 |
| Item Lure ERS* | 0.74 | 2.93 | 0.005 | Item Lure ERS* | 0.74 | 2.9 | 0.006 |
| Theme Lure ERS* | -0.5 | -2.16 | 0.04 | Theme Lure ERS | -0.48 | -1.87 | 0.07 |
| Target ERS x Age | -0.007 | -0.02 | 0.98 | Target ERS x Age | 0.002 | 0.006 | 0.99 |
| Item Lure ERS x Age | -0.28 | -0.81 | 0.42 | Item Lure ERS x Age | -0.28 | -0.79 | 0.44 |
| Theme Lure ERS x Age | 0.21 | 0.65 | 0.52 | Theme Lure ERS x Age | 0.19 | 0.59 | 0.56 |
|  |  |  |  | Global ERS | -0.02 | -0.24 | 0.81 |
| ROI: Lateral Occipital Cortex |  |  |  | ROI: Lateral Occipital Cortex |  |  |  |
| Intercept* | 0.42 | 7.29 | <.001 | Intercept* | 0.41 | 6.4 | <.001 |
| Age* | -0.28 | -3.42 | 0.001 | Age* | -0.28 | -3.92 | 0.001 |
| Target ERS | 0.04 | 0.141 | 0.89 | Target ERS | 0.03 | 0.11 | 0.91 |
| Item Lure ERS | -0.29 | -1.21 | 0.23 | Item Lure ERS | -0.31 | -1.25 | 0.22 |
| Theme Lure ERS | -0.03 | -0.12 | 0.91 | Theme Lure ERS | -0.06 | -0.2 | 0.84 |
| Target ERS x Age | -0.15 | -0.43 | 0.67 | Target ERS x Age | -0.13 | -0.36 | 0.72 |
| Item Lure ERS x Age* | 0.71 | 2.28 | 0.03 | Item Lure ERS x Age* | 0.7 | 2.21 | 0.03 |
| Theme Lure ERS x Age | -0.3 | -0.89 | 0.39 | Theme Lure ERS x Age | -0.31 | -0.9 | 0.37 |
|  |  |  |  | Global ERS | 0.03 | 0.39 | 0.69 |
| ROI: Posterior Hippocampus |  |  |  | ROI: Posterior Hippocampus |  |  |  |
| Intercept* | 0.29 | 10.49 | <.001 | Intercept* | 0.29 | 10.29 | <.001 |
| Age* | -0.15 | -3.58 | <.001 | Age* | -0.15 | -3.45 | 0.001 |
| Target ERS | 0.3 | 0.75 | 0.46 | Target ERS | 0.28 | 0.66 | 0.51 |
| Item Lure ERS | -0.17 | -0.57 | 0.57 | Item Lure ERS | -0.19 | -0.58 | 0.56 |
| Theme Lure ERS* | -0.69 | -2.06 | 0.04 | Theme Lure ERS | -0.71 | -1.95 | 0.06 |
| Target ERS x Age | -0.34 | -0.71 | 0.48 | Target ERS x Age | -0.33 | -0.67 | 0.51 |
| Item Lure ERS x Age | 0.36 | 0.91 | 0.37 | Item Lure ERS x Age | 0.37 | 0.9 | 0.37 |
| Theme Lure ERS x Age | 0.57 | 1.29 | 0.21 | Theme Lure ERS x Age | 0.57 | 1.27 | 0.21 |
|  |  |  |  | Global ERS | 0.02 | 0.19 | 0.88 |

\* indicates $p < .05$

We also examined the influence of global ERS in predicting false memory rates. To do so we repeated the above multiple linear regressions while including global ERS as a predictor (see Table 2). When accounting for the global ERS signal, item lure ERS still positively predicted false memory rate in early visual cortex (b(39) = 0.75, *p*=.006), while the effect of thematic lure ERS approached significance (b(39) = −0.48, *p*=.06). In lateral occipital cortex, the significant age group X item lure ERS interaction also remained significant (b(39) = 0.70, *p*=.03), such that item lure ERS negatively predicted false memory rates in older adults but not younger adults. Finally, in posterior hippocampus, thematic lure ERS marginally predicted false memory rates (b(39) = −0.71, *p*=.06). Critically, however, global ERS did not significantly predict false memory rates in any of aforementioned regions (all *p’s* > .05).

Like above, we reconducted the above analyses while conducting permutation testing to confirm the significant findings and correct for the effect of false positives. Specifically, we repeatedly shuffled the data randomly and reconducted our multiple linear regression analyses (Winkler et al., 2014). We repeated this procedure 10,000 times for each regression to build a null distribution to simulate the potential relationship between false memories and ERS if our effects did not exist. We found no differences in significance after conducting permutation testing.

## 4. Discussion

The current study investigated potential age differences in ERS associated with memory for targets, item lures, and thematic lures, and how these age differences contribute to false memory rates in younger and older adults. Our results showed an overall age deficit in ERS across all three trial types of interest within ROIs in the visual system and portions of the hippocampus. We also observed that ERS within these regions was behaviorally relevant to false memory differentially in younger and older adults. Specifically, ERS within early visual cortex and posterior hippocampus contributed false memory rates across the entirety of our sample, and ERS in lateral occipital cortex predicted within older adults alone. Finally, we found that global ERS, as measured by neural similarity between all items at retrieval and encoding, accounted for age deficits in single-item ERS in occipital and hippocampal regions, but could not account for the relationship between ERS and false memories in aging. Rather, only single-item ERS contributed to false memory rates. We discuss these findings further below.

### 4.1 Age deficits in ERS

Recent work suggests that the similarity between patterns of neural activity across memory phases is a critical indictor of both veridical and false memories (Ritchey et al., 2013; Wheeler & Buckner, 2004; Ye et al., 2016). Such similarity is thought to reflect the recapitulation of sensory signals related to the studied items and components of memory processing such as basic perceptual features including item color, orientation, and size. It is theorized that greater recapitulation of this information between encoding and retrieval will result in higher rates of true, or false, memories depending on the stimulus or memory signal being recapitulated. It is also speculated that age differences in this recapitulation are an underlying mechanism contributing to age-related memory impairments (Brainerd & Reyna, 2002; Dennis & Turney, 2018). Our findings showed widespread age deficits in ERS throughout the ventral visual cortex and hippocampus. Specifically, ROIs within bilateral early visual cortex, lateral occipital cortex, fusiform gyrus, and posterior hippocampus showed reduced similarity of neural patterns of activation across memory phases in older compared to younger adults. Critically, this reduction was associated not only with the need to identify previously viewed stimuli as “old” (target-target ERS), but also the need to recapitulate target related details in order to correctly identify novel stimuli as “new” (target-item lure and target-theme lure ERS).

The notion of sensory reactivation in memory is a long-standing theory (Damasio, 1989; James & Burkhardt, 1983; Kosslyn, 1994) that is supported by univariate findings showing that neural activity is greater at retrieval for remembered compared to forgotten memories (Baym & Gonsalves, 2010; Vaidya et al., 2002; Wheeler et al., 2000). Additionally, a consistent finding in aging and memory research is that older, compared to younger, adults exhibit reduced activity in visual cortices associated with such memory success (Dennis et al., 2008; Devitt & Schacter, 2016; Gutchess & Schacter, 2012). These age-related reductions in neural activity within the visual cortex, coupled with age-deficits in recollection (Dennis et al., 2012; McCabe et al., 2009) has led to the conclusion that older adults do not retrieve sensory details related to the original memory episode as often or as vividly as do young adults. The observed age-related reduction in ERS within both the occipital cortex and hippocampus supports and extends this corpus of univariate findings with greater specificity. Specifically, our results suggest that, compared to younger adults, older adults exhibit a significant reduction in the recapitulation of neural activation patterns between the encoding of a visual stimulus and the retrieval of the same or related stimulus. That is, age reductions in ERS were found not only for target recognition, but also associated with item and category lure decisions.

While ERS does not provide details about what information, specifically, is being processed at encoding and retrieval, age-related reductions in ERS may be suggestive of a number of underlying information processing deficits in older adults. Reduced neural recapitulation in sensory cortices is likely a reflection of impaired reactivation of perceptual details, as predicted by the sensory reactivation hypothesis (Slotnick & Schacter, 2004). For example, it is possible that older adults do not store, and thus do not have available as many or as high-quality, encoding-related details for retrieval compared to their younger counterparts (Dennis, Bowman, et al., 2014; Hill et al., 2021). It is also possible that older adults have such details stored in their memory, but fail to reactivate these memory traces at the time of retrieval (Trelle et al., 2020; Naspi et al., 2021). Either possibility would result in overall reduce similarity between neural patterns within occipital cortices across encoding and retrieval. Recent work from Hill et al. (2021) provides evidence for the former explanation, as age deficits in neural reinstatement became nonsignificant after controlling for the distinctiveness of neural patterns in visual regions during encoding. Similarly, age deficits in single-item ERS within the current study became nonsignificant after controlling for global ERS capturing the neural pattern similarity of all items viewed at encoding (see below for more discussion on global ERS deficits and relation to behavior). Such results suggest that older adults exhibit reduced pattern similarity at multiple levels, and potentially at multiple stages of memory processing. Future work should examine how reduced ERS in older adults is reflective of retrieval processing failures.

Interestingly, within the current results was the finding that the age-reduction in ERS was consistent across both target and lure conditions. While the need to retrieve encoding-related details and recapitulate encoding processes in support of veridical memory recognition is important, veridical memory endorsement in recognition tasks can be based on retrieval of gist-level memory traces (Brainerd & Reyna, 2002), absent of retrieval for fine-grain details of past experiences. Thus, reduced ERS within regions supporting detailed perceptual representations of past episodes may not be a particular detriment to veridical retrieval if more general properties of the encoding episode are reinstated at retrieval. This is reflected in our lack of age deficits for target memory (i.e., hit rate). Yet, while accurate target identification can be supported in the absence of detail-level reinstatement, the accurate rejection of a perceptually related lure cannot. That is, if a novel object shares overlapping gist information with a target item, as in the case of perceptually related item-lures, there is a direct benefit to lure identification with recapitulation of item-related details. Age-related reductions in ERS between encoded targets and the presentation of perceptual lures at retrieval would contribute to higher false memory rates in older adults, especially if there was no reduction in ERS of gist representations, or if older adults do not re-engage the same neural processes.

Regarding behavior, older adults who had greater item lure ERS in lateral occipital cortex exhibited lower rates of false memories. It is likely that older adults who are able to recapitulate the gist signal of an item lure are more likely to correctly identify and reject perceptual lure items. Specifically, information between encoding and retrieval may become coarser with age, or more gist-like, but recapitulation of this information benefits processes associated with rejecting the lure item. Previous univariate neuroimaging studies have implicated late visual regions in conscious remembering and semantic labeling of lure items (Slotnick & Schacter, 2004). In the current study, older adults who recapitulated more semanticized or gist-like information were more likely to correctly reject the perceptual lure. This is supported by previous univariate neuroimaging work from our group finding older adults preferentially recruit late visual cortical regions when correctly rejecting lure items (Bowman & Dennis, 2015). It is possible that a shift occurs in healthy aging regarding the content of information recapitulated between encoding and retrieval, where recapitulation of neural patterns within late visual regions becomes relevant for rejecting perceptual lure items within older adults (Bowman et al., 2019). Contrary to the negative relationship between ERS and false memories within lateral occipital cortex, we also observed that individuals who exhibited greater item lure ERS in the early visual cortex also tended to exhibit increased rates of false memories, regardless of age. Early visual regions have been found to be associated with recapitulating sensory signals during retrieval (Buckner & Wheeler, 2001; Rugg & Wilding, 2000; Vaidya et al., 2002). While the overall strength of recapitulation of lure items was reduced by age, it appears the content of information associated with the lure items is preserved in aging within early visual cortex (Koen & Rugg, 2019). That is, participants who generally recapitulate visual information associated with the encoded item to a greater degree in the presence of the perceptual lure are more likely to falsely endorse the lure item (Slotnick & Schacter, 2006). Collectively, such findings highlight the subtle role of how regions within visual cortex differentially contribute to false memories, and how ERS within such visual regions is impacted by age.

In addition to age-related reductions in ERS within the visual cortex, reductions were also observed within portions of the hippocampus. Notably, we found an overall age deficit in ERS in posterior hippocampus. This finding is consistent with previous work by Abdulrahman et al. (2017), who interpreted reduced reinstatement effects in the hippocampi of older compared to younger adults as reflective of dedifferentiation of neural patterns associated with pattern separation and completion processes. Furthermore, within this region, thematic lure ERS negatively contributed to false memories across both younger and older adults within our sample (although this finding became marginal after controlling for global ERS). A large body of literature implicates the hippocampus, and specifically dentate gyrus within the hippocampus, in pattern separation processes (Brock Kirwan et al., 2012; Yassa et al., 2011; Yassa & Stark, 2011). Here we find that although older adults exhibit reduced ERS in posterior hippocampus across all memory conditions, participants with increased thematic lure ERS demonstrated more correct rejections. This is likely indicative of semantic feature processing utilized for pattern separation, as research in young adults suggest participants often reject lure items of greater conceptual (as opposed to perceptual) confusability in a mnemonic similarity task (Naspi et al., 2021).

The finding of this relationship between ERS and ability to reject related lures builds upon recent work showing that portions of the hippocampus exhibit differential activity when correctly identifying lures as “new” as compared to correctly identifying targets as “old” during memory retrieval in younger adults (Klippenstein et al., 2020). Specifically, Kleippenstein et al. (2020) observed greater neural activation in the hippocampal head and body for correctly identifying targets as “old” compared to identifying lures as “new”. That is, combined with previous univariate work examining false memories (Bowman et al., 2017; Bowman & Dennis, 2015), the current results suggest that that not only is the amount BOLD activity relevant to the correct rejections of lures, but that the degree of recapitulation of the pattern of neural activation exhibited between encoding and retrieval also facilitates the correct identification of lures as novel items. Future research should consider how ERS varies within subfields the hippocampus, and how such recapitulation might relate to pattern separation processes in younger and older adults.

Interestingly, we failed to observe any age deficits in ERS, or a relationship between ERS and false memories, in regions outside of the visual stream or hippocampus. While past research in false memories has implicated a strong role in PFC and parietal cortex underlying the endorsement of false memories at retrieval (Atkins & Reuter-Lorenz, 2011; Garoff-Eaton et al., 2006; Kim & Cabeza, 2007; Kurkela & Dennis, 2016), the current results do not find that this retrieval-related processing is linked with processing at the time of encoding. Such findings are consistent with a recent report by Trelle et al (2020), who found that item-level reinstatement was not reduced by age in angular gyrus. It appears that while visual information contained in memory traces recapitulated between encoding and retrieval are reduced by age, downstream regions are able to maintain the same correspondence of neural patterns between memory phases in the context of attentional or higher-order processes. Further, such regions have been extensively implicated in retrieval processes associated with false recollection (Atkins & Reuter-Lorenz, 2011; Kim & Cabeza, 2007). While fronto-parietal regions are involved in monitoring of lure information, such functions differ from their role during encoding. This discrepancy between encoding and retrieval could result in the lack of any age or trial effects, or relationships with behavior. Future work examining pattern similarity in fronto-parietal regions during encoding and retrieval separately may help resolve this discrepancy.

### 4.2 Global ERS

In addition to calculating single-item ERS, we also calculated a global ERS metric within each ROI to account for group differences in baseline similarity between encoding and retrieval in our analyses (Ye et al., 2016). As a follow up to the single-item ERS analysis, we found that differences in global ERS accounted for observed age deficits in ERS in the visual cortex and posterior hippocampus, but did not contribute to behavioral measures of false memory rates in any region. Such findings suggest that age deficits in ERS occur at multiple levels, including the single-item (i.e., “the similarity between a specific dog at retrieval and a specific dog at encoding”) as well as a more holistic (i.e., “the similarity between everything one is seeing at retrieval and everything one has seen at encoding”) level. It is also possible that age deficits in ERS also occur at an intermediate category or gist-level. While the current design precludes such an analysis, future work could examine at just what levels such age-related deficits occur. Further, this information may be useful to investigators who wish to investigate age-related reductions in ERS, but whose designs do not allow for the examination of ERS at the single-item level, such as when a lure at retrieval is related to multiple exemplars viewed during encoding. Finally, with regard to the effect or global ERS on item-level ERS, a recent report observed a similar effect when examining neural distinctiveness during encoding at the single-item and global level (Kobelt et al., 2021). While older adults exhibited reduced neural distinctiveness at both the single-item and global level, only neural distinctiveness of the item level predicted veridical memory rates (Kobelt et al., 2021). Taken together, results highlight the importance of item-level ERS in explaining the age-related increase in false memory rates in older compared to younger adults.

As noted above, while global ERS was able to account for variance in age differences with regard to single-item ERS, global ERS did not fully account for the relationship between ERS and behavior. That is, even when including global ERS as a covariate in the regression models, it was the single-item ERS that contributed to false memory rates in lateral occipital cortex and early visual cortex. Such findings highlight how direct correspondence of neural patterns associated with single items (i.e., a specific stimulus at encoding and its corresponding stimulus at retrieval) is predictive of false memory above and beyond a holistic measure of neural correspondence between memory phases. While single-item ERS in posterior hippocampus was predictive of false memory rates, this became marginally significant after accounting for baseline differences global ERS. This may be due to a lack of power due to a relatively small sample size. Future studies should therefore employ larger samples to examine the influence of age on the correspondence between ERS and behavior.

### 4.3 Trial type effects

Irrespective of age, we observed trial type effects within our ERS analysis throughout the visual system and portions of the medial temporal lobe. Specifically, ERS was greater for targets compared to either lure condition in lateral occipital cortex. This difference is likely reflective of greater sensory recapitulation in response to previously seen items when that item is being viewed than when any related item is presented (Slotnick et al., 2012). Such greater ERS could be triggered by the actual details inherent within the target item as opposed the absence of such details in lure items. In contrast to greater target ERS in sensory regions, comparatively greater ERS for item lures was observed in MTL regions including posterior hippocampus and entorhinal cortex. While counterintuitive, this pattern of ERS could be reflective of neural responses to specific cues within the lure item that trigger the need to retrieve details of the target item. Specifically, Robin & Moscovitch (2017) hypothesize that posterior hippocampus and entorhinal cortex are involved collaboratively in elaboration processes during retrieval in response to cues. Relatedly, patient lesion studies suggest that entorhinal cortex is involved in familiarity processes associated with false memory (Brandt et al., 2018). In conjunction with work by Robin & Moscovitch, participants potentially recapitulate neural patterns associated with familiarity when presented with specific perceptual cues (i.e., a dog similar to one viewed previously), and that such information is modulated within entorhinal cortex. Finally, regarding greater item lure ERS in posterior hippocampus, a recent meta-analysis suggests that posterior hippocampus is preferentially activated while integrating fine-grained representations as opposed to coarse-grained representations during episodic retrieval (Grady, 2020). Perceptually similar stimuli, such as the item lures used in the current study, likely require recapitulation of fine-grained information within posterior hippocampus to assist in pattern separation processes. Future work with high-resolution neuroimaging will assist in understanding how the components of the hippocampus and medial temporal lobe differentially recapitulate neural patterns associated with false recognition.

### 4.4 Limitations and Future Directions

The current study has several limitations that should be considered. First, our study is correlational in nature so we cannot claim increased ERS for lure items causes increases in false memories, only that the two constructs are related. Second, by using a cross sectional sample we cannot examine age-related changes in ERS, only differences between younger and older adults. Longitudinal studies will assist in elucidating how the relationship between ERS and memory errors is altered by increasing age. Third, due to the of current study design, every item viewed during encoding is its own perceptual category. We are therefore unable to calculate neural distinctiveness during encoding to use as a baseline neural similarity measure, as recent studies have with face and house categories of visual stimuli (Hill et al., 2021; Kobelt et al., 2021). However, we contend that global ERS is an appropriate and theoretically meaningful baseline assessment of recapitulation between younger and older adults when assessing single-item ERS. Regarding future directions, recent work suggests that applying excitatory transcranial magnetic stimulation to dorsolateral prefrontal cortex increases pattern similarity in visual cortex during encoding and retrieval, as well as single-item ERS in the hippocampus in younger adults during an episodic memory task (Wang et al., 2018). An exciting avenue for future research is therefore to examine whether such TMS application increases ERS in older adults, and if ERS increases are associated with alterations in false memory rates. Furthermore, it remains to be seen how cognitive training regimens may alter ERS in older adults alongside beneficial changes in memory performance.

## 5. Conclusions

In conclusion, the current study investigated age-related deficits of ERS associated with target and lure items, and how such deficits relate to false memory rates. We observed age-related reductions in ERS throughout the ventral visual stream as well as in portions of the hippocampus. Age deficits in global ERS accounted for single-item ERS age reductions, however only single-item ERS predicted individual differences in false memory rates. Such results carry implications regarding age deficits of neural patterns across memory phases, and how such neural similarity is related to memory errors of commission. Specifically, they suggest recapitulation of lure information within occipital regions is utilized to both false alarm to, and correctly reject, previously unseen items. Future longitudinal and intervention work will assist in deepening our understanding regarding how ERS changes with age, and how to ameliorate age-related memory errors in older adults.

## Acknowledgements

We thank the Penn State Social, Life, & Engineering Sciences Imaging Center (SLEIC), 3T MRI Facility. This work was supported by a National Science Foundation grant BCS1025709 and BCS2000047 awarded to NAD. It was also supported by dissertation awards granted by the American Psychological Association and Penn State’s Research and Graduate Studies Office to CRB.

